# Context information supports serial dependence of multiple visual objects across memory episodes

**DOI:** 10.1101/667626

**Authors:** Cora Fischer, Stefan Czoschke, Benjamin Peters, Benjamin Rahm, Jochen Kaiser, Christoph Bledowski

## Abstract

Visual perception operates in an object-based manner, by integrating associated features via attention. Working memory allows a flexible access to a limited number of currently relevant objects, even when they are occluded or physically no longer present. Recently, it has been shown that we compensate for small changes of an object’s feature over memory episodes, which can support its perceptual stability. This phenomenon was termed ‘serial dependence’ and has mostly been studied in situations that comprised only a single relevant object. However, since we are typically confronted with situations where several objects have to be perceived and held in working memory, the central question of how we selectively create temporal stability of several objects has remained unsolved. As different objects can be distinguished by their accompanying context features, like their color or temporal position, we tested whether serial dependence is supported by the congruence of context features across memory episodes. Specifically, we asked participants to remember the motion directions of two sequentially presented colored dot fields per trial. At the end of a trial one motion direction was cued for continuous report either by its color (Experiment 1) or serial position (Experiment 2). We observed serial dependence, i.e., an attractive bias of currently toward previously memorized objects, between current and past motion directions that was clearly enhanced when items had the same color or serial position across trials. This bias was particularly pronounced for the context feature that was used for cueing and for the target of the previous trial. Together, these findings demonstrate that coding of current object representations depends on previous representations, especially when they share similar content and context features. Apparently the binding of content and context features is not completely erased after a memory episode, but it is carried over to subsequent episodes. As this reflects temporal dependencies in natural settings, the present findings reveal a mechanism that integrates corresponding bundles of content and context features to support stable representations of individualized objects over time.

---

Visual cognition relies heavily on the interplay between perception and memory processes. Behavioral research has shown that we create distinguishable visual objects from a current scene by integrating individual features that belong together (Treisman, 1986). Moreover, we are able to maintain mental representations of these objects in working memory (WM) over short periods of time (Luck & Vogel, 1997). This enables us to use object information flexibly for a variety of cognitive tasks even when they are occluded or physically no longer present. However, which objects need to be maintained in WM can change from one moment to another, because the world around us changes constantly. As situations often change gradually instead of abruptly, many of these changes are foreseeable, and current object representations can often be based on preceding ones. Thus, the exploitation of such short-term dependencies over time represents an important requisite of perceived environmental stability.

A series of recent studies have examined in detail how an object representation that is currently encoded into WM is influenced by an object representation that was encoded in the previous trial. In the seminal study by Fischer and Whitney (2014), participants encoded the orientation of a Gabor patch. After a short delay, they were asked to report it in a continuous manner, i.e. by orienting a response line to match the memorized orientation. They found that the reported orientation in the current trial was systematically attracted by the orientation remembered in the previous trial. Importantly, this attractive bias was strongest for orientations that were about 30° apart, and decreased rapidly for larger orientation differences, which shows that it was tuned to the feature similarity between objects across trials. Additional experiments showed that the bias was enhanced by spatial and temporal item proximity. Furthermore, they observed that items needed to be attended and encoded in order to elicit this bias on subsequent items. Fischer and Whitney (2014) coined the term ‘serial dependence’ for this attractive bias, as it shows a systematic and highly specific dependence of serially encoded objects. Serial dependence has since been observed for other stimulus types, such as faces (Liberman, Fischer & Whitney, 2014), spatial positions (Bliss, Sun & D’Esposito, 2017) or ensemble representations (Manassi, Liberman, Chaney & Whitney, 2017) (see Kiyonaga, Scimeca, Bliss & Whitney, 2017 for an overview). As serial dependence reduces the perceptual difference between consecutive objects that are similar to each other, it has been interpreted as a mechanism that promotes perceptual stability and continuity of a visual object over time (Fischer & Whitney, 2014).

Most studies so far have examined a special case that occurs rarely in nature, i.e. they studied serial dependence in a situation that comprised only a single relevant object. In the real world, however, we are commonly confronted with situations where several objects have to be perceived and maintained in WM. The central question of how we create stability of several object representations over time has remained unsolved.

A possible answer to this question refers to the idea that objects are maintained in WM as integrated representations of their features (Luck & Vogel, 1997; Treisman, 1986). Such multi-feature representations would facilitate associations between objects over time. Alternatively, multi-feature objects could be represented in WM by a simultaneous, but independent storage of individual features (Bays, Wu & Husain, 2011). Reconciling both approaches, Brady, Konkle and Alvarez (2011) have proposed that information in WM is structured as a hierarchical feature bundle consisting of two levels. The top level of a bundle represents an integrated object, while the bottom level contains low-level features that are stored independently. In line with this model, Oberauer and Lin (2017) have proposed that objects in WM consist of several features that are bound together. They explicitly distinguish between object ‘content’ denoting the feature that needs to be reported, and object ‘context’ representing the feature dimensions via which an object can be cued for report. Context feature can hence refer to the spatial position or the serial position in an encoding sequence, but also to other object features like color. Moreover, context features can differ with regard to their relevance for the ongoing cognitive task. Context features that serve as cues to identify the currently task-relevant object can be referred to as ‘task-relevant context features’. In contrast, context features that also differ between objects but are neither reported nor serve as a cue can be referred to as ‘task-irrelevant context features’ (Figure 1a).

**Figure 1.**
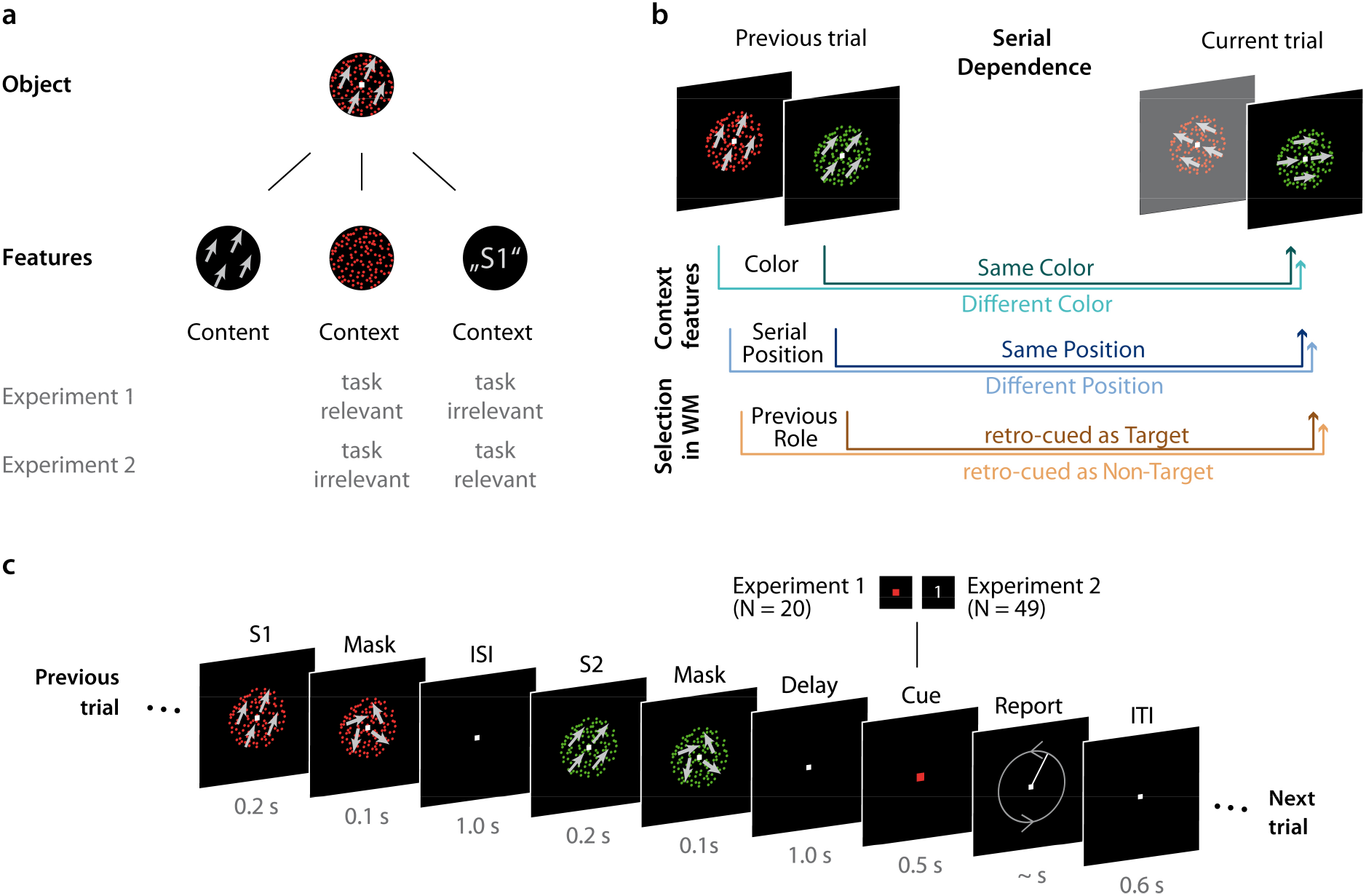
Object definition and experimental paradigm. a) Every object can be defined regarding its content feature, here motion direction of 25°, and regarding its context features, here color (red) and serial position (first in the sequence, S1). The context feature by which targets were cued was the task-relevant feature, whereas the other feature was task-irrelevant (e.g. color and serial position in Experiment 1, respectively). b) We assessed the effects of three factors on the response error for a cued item (target) in the current trial: color (same or different), serial position and role (target or non-target) of an item in the previous trial. c) In every trial, participants had to memorize motion directions of two sequentially presented dot fields (S1 and S2) and report one of them after a short delay by adjusting the orientation of a line. The to-be-reported target item was cued via color (Experiment 1) or via serial position (Experiment 2).

The present study investigated whether multiple objects that are encoded into WM can be related to each other across memory episodes via corresponding context features. If this was the case it would indicate that not only content features, but also context features leave traces in memory that support serial dependence between objects. Furthermore, as the task-relevant context should at least be attended more strongly than the task-irrelevant context, the former should have a stronger impact on serial dependence than the latter. Finally, serial dependence could also be influenced by whether or not a previous object served as a target that was selected and reported (Czoschke, Fischer, Beitner, Kaiser & Bledowski, 2019). As attentional selection and retrieval processes affect the representations themselves (Griffin & Nobre, 2003; Landman, Spekreijse & Lamme, 2003; Souza & Oberauer, 2016), they are likely to influence the strength of the traces objects leave behind in WM, too. Accordingly, targets should produce a stronger serial dependence.

To answer these questions, we conducted two experiments, in which two motion directions per trial were sequentially presented as colored dot fields. After a short delay, one motion direction was retrocued and had to be reported by adjusting the direction of a probe line. In the first experiment, color was the task-relevant context feature by which the target was cued. In the second experiment, the item was cued by its serial position. This enabled us to investigate the impact of three factors that may promote serial dependence of multiple visual objects across memory episodes: the congruence of context features, their task-relevance, and internal attentional selection of a target object (Figure 1b).

## Experiment 1: Color Cueing

### Methods

#### Subjects

Twenty-two subjects who were recruited from the Goethe-University Frankfurt and the Fresenius University of Applied Sciences Frankfurt participated in experiment 1. All subjects reported normal or corrected-to-normal vision. Two subjects aborted the experiment after the practice trials due to difficulties to perform the task. This resulted in a sample of 20 subjects (10 male), aged between 18 and 32 years (mean: 23.3 years). All subjects gave informed consent and were compensated with € 10/h or course credit. The study was approved by the Ethics Committee of the Medical Faculty of the Goethe-University Frankfurt am Main and therefore complied with their ethical regulations.

#### Stimuli

Random dot patterns (RDP) were presented centrally on the screen and consisted of 200 dots colored in red (RGB: [255, 0, 0]) or green (RGB: [0, 0, 255]) on a black (RGB: [0, 0, 0]) background. The dots were presented within an invisible circular aperture which had a radius of 7.5° of visual angle. The dots had a diameter of 0.15° of visual angle and were placed randomly within the circular aperture of the RDP at stimulus onset. The dots moved with a velocity of 3.5°/s and fully coherent in a direction randomly drawn from a pool of directions between 5° and 355° spaced 10° from one another, therefore avoiding cardinal directions. Dots reaching the edge of the aperture were repositioned randomly on the edge of the opposing side of the aperture, so that dot density was kept constant throughout the presentation. Throughout the whole experiment a white fixation square with a diagonal of 0.15° of visual angle was presented centrally on the screen, except for the cue presentation, when the fixation square changed its color to red or green to cue which item should be reported. The item was reported by adjusting a randomly oriented line to match the recalled direction. The response line was white, with a width of 0.6° and a length equaling the dot field radius. The starting point of the line was the fixation square and the end point could be altered so that the line could point in all possible directions.

#### Procedure

Experiment 1 consisted of a delayed-estimation task, in which two sequentially presented motion directions had to be retained in memory, one of which after a short delay was cued for report (Figure 1c). Specifically, subjects saw two sequentially presented RDPs per trial (S1 and S2), each with a different motion direction and a different color, either red or green. Each trial began with the presentation of the first stimulus (S1) for 200 ms followed by a noise mask for 100 ms consisting of dots moving with 0% coherence (i.e., randomly) and of the same color as the preceding RDP. After a 1000 ms interval (ISI) the second stimulus (S2) and its noise mask were presented for 200 ms and 100 ms, respectively. Following a delay of 1000 ms, the fixation square changed its color to red or green for 500 ms, thereby cueing which motion direction had to be reported. Immediately after cue offset, a randomly oriented line was presented. Subjects had to report the motion direction by rotating the line via horizontal mouse movements. No time limit was given for the response and subjects were encouraged to work as precisely as possible. After adjusting the line direction, subjects had to confirm their response by pressing the left mouse button. If the entered direction differed more than 30° from the cued direction, another line pointing in the correct direction was presented for 500 ms as an error feedback (see Kang & Choi, 2015, for a similar procedure). At the end of each trial, a fixation screen of 600 ms was presented. Subjects were instructed to fixate the fixation square throughout the whole experiment.

In every trial, two different directions were presented, differing between 10° and 170°, equally spaced in steps of 10°, from one another. Over trials, the direction differences were balanced so that every possible direction difference occurred equally often. The order of the dot field colors was balanced across trials, so that both items were presented in both colors equally often, but in each trial the two stimuli had different colors from one another. In half of the trials, the red stimulus had to be reported, and in the other half the green one, which was balanced over encoding positions and direction difference combinations. The order of the trials was not balanced, therefore resulting in a different number of trials per subject and condition for the serial dependence analysis. Each item could be described with regard to its content (i.e., its motion direction), its context (i.e., its color and serial position) and its previous role (i.e., whether it was a target or a non-target) (see Figure 1a). Additionally, here the context feature color served as the task-relevant cueing feature, whereas the context feature serial position was task-irrelevant (see Experiment 2 below for the opposite assignment). The bias produced by an item from a previous trial on the report of the target item of the current trial could therefore be investigated with regard to three analysis factors with two levels each: item color (same or different color as the target of the next trial), serial position (same or different serial position as the target of the next trial) and previous role (target or non-target) (Figure 1c). Therefore, the impact of three factors on serial dependence was examined in a 2*2*2 design. On average, 392.24 trials per factor level combination were analyzed in Experiment 1, with an average 10.6 trials per factor level combination and distance, and 393.46 and 10.6 trials in Experiment 2, respectively.

Every subject completed 1632 trials in two sessions on different days, lasting for approximately 2 hours each (including instruction and practice trials). Each session was divided into eight blocks of 102 trials with self-paced breaks in between. After 45 minutes a general break was given, which was around the half of one session for most subjects. Up to three subjects completed the experiment in parallel in a dimly lit room, acoustically and visually shielded from one another. Subjects were seated at a viewing distance of approximately 50 cm from the display. MATLAB software with the Psychophysics Toolbox extensions (Brainard, 1997; Pelli, 1997) was used for stimulus generation and presentation. Three different LCD-monitors with a 4:3 display ratio and running with 60 Hz refresh rate were used.

#### Analysis

Prior to the estimation of serial dependence, we excluded trials in which the response error was at least 3 SDs higher than the subject’s mean response error, or in which the response time exceeded 20 s, indicating potential attentional lapses, accounting on average for 2.74% of trials in Experiment 1 and 2.47% in Experiment 2. We also excluded the first trial of each session as well as trials following a break, because in the analysis of serial dependence we were interested in the effect of the previous trial on the current one. We also demeaned the response errors by subtracting the overall mean response error of a participant from each individual response error to remove general individual response biases independent of serial dependence.

The evaluation of serial dependence was based on individual response errors, defined as the deviation between presented and entered direction. The errors were sorted regarding the difference between the target stimulus of the current trial and a stimulus from the previous trial as well as the relation of difference between the current item and the item of the previous trial (clockwise or counter-clockwise). For the target item of a current trial, the influences of two different items of the previous trial (S1 previous trial and S2 previous trial) were evaluated separately. The difference was computed by subtracting the direction of the current item from the direction of the item of the previous trial. Therefore, when the current item was oriented more clockwise or more counter-clockwise, this resulted in a negatively or positively signed distance, respectively. A mean response error for a signed distance (distance*relation) deviating from 0 indicated a systematic response bias. When the sign of this systemic bias matched the sign of the distance between the directions, it indicated an attractive response bias. Conversely, an opposite sign of the systematic bias compared to the signed distance indicated a repulsive response bias.

Trials were then sorted according to their respective levels of the three analysis factors (see Procedure) before computing the mean response bias per signed distance for each factor level combination. Then we could examine the effects of the two context factors as well as task relevance on serial dependence (see Fig. 1C). To assess the effect of those factors, we computed the mean response bias per factor level by averaging response biases of the appropriate conditions. For example, to examine the effect of color, we computed the mean of mean response biases of all conditions where items had the same color (regardless of serial position and task relevance) and different colors, which resulted in one mean response bias per level of the factor “color”, signed distance and subject.

The individual mean response biases were used to evaluate the serial dependence per contrast level. We fitted the first derivative of a Gaussian curve (DoG; e.g. Fischer & Whitney, 2014), a model which is usually used to describe serial dependence. The DoG, given by

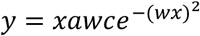

was fitted to the pooled mean response biases of all subjects (similar to the procedure by Fritsche et al., 2017) per factor level, i.e. one data point per subject and distance for the respective factor level. In the DoG, x is the relative direction difference of two stimuli, a is the amplitude of the curve peak, w scales the curve width, and c is the constant 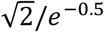. The w parameter was constrained to a value range of .01 to .1. We optimized the log likelihoods of our curve fitting using Bayesian adaptive direct search (BADS; Acerbi & Ma, 2017). BADS alternates between a series of fast, local Bayesian optimization steps and a systematic, slower exploration of a mesh grid. To estimate the variability of the parameters a and w, we bootstrapped the DoG curve fit 1,000 times, sampling the data with replacement on each iteration, and computed the standard deviation of the resulting bootstrapped distributions of a and w (see Fischer & Whitney, 2014, for a similar procedure). For an easier interpretation of the w parameter, we converted it into the full width at half maximum (FWHM).

#### Statistical analysis

Permutation tests were used to assess effects on the group level. Specifically, to test whether there was a significant serial dependence, we randomly inverted the signs of each participant’s mean response error. Subsequently, we fitted a new DoG model to the pooled group data and collected the resulting amplitude parameters a in a permutation distribution. We repeated this permutation procedure 1,000 times. As p-values we report the percentage of permutations that led to equal or higher values for a than the one estimated for the empirical data. Based on previous findings (Czoschke et al., 2018; Fischer & Whitney, 2014; Fritsche et al., 2017), we expected a positive serial dependence and therefore the significance level was set to α = 0.05 (one-sided permutation test). Significance of the model fit was assessed by a permutation test of the a parameter only. The w parameter was constrained to a range that excludes zero and zero would be the expected value of w if the fitted data randomly fluctuated around zero without a systematic bias. To test whether serial dependence differed significantly between factor levels, we applied the same procedure as above except that we randomly shuffled the labels of factor levels per participant. Thereby we generated a distribution of a and w differences, against which the a and w difference of the empirical data could be tested. Based on our hypotheses, we expected an enhancement of serial dependence, either in strength (i.e., higher amplitude) or width (i.e., broader tuning width) for same color or serial position in comparison to different color or serial position as well as for targets in comparison to non-targets. Therefore, the significance level was set to α = 0.05 (one-sided permutation test). The p-value was given by the proportion of permuted differences whose values were equal or greater than the empirical difference.

The same permutation procedure was used to examine possible interactions between the three investigated factors. To this aim, we investigated the difference of differences for two factors. For example, to investigate an interaction between task relevance and color, we computed the difference same vs. different color separately for targets and non-targets, which resulted in two differences. As an indicator for an interaction, we computed the difference between those two differences. In the permutation procedure, in each iteration (1,000) the labels of factor levels for both factors were shuffled per participant. As we did not have clear expectations regarding interactions between the factors, we conducted a two-sided permutation test, i.e. the p-value was given by the proportion of permuted differences, whose absolute values were equal or greater than the absolute empirical difference. The significance level was set to α = 0.05.

#### Effect size calculation

Our fitting procedure yielded group estimates for amplitude and width, whereas for the calculation of mean-based effect sizes, individual estimates are necessary. As there is no standard procedure for estimating the effect sizes for our analysis, we aimed at obtaining an approximation of the effect sizes that reflects the effects revealed by our permutation tests as good as possible. An estimation of the effect sizes analog to Cohen’s d was calculated separately for the amplitude and width of the serial dependence, based on the obtained fittings for every condition. We used computations of the individual data informed by the group fitting to obtain individual estimates. Specifically, for amplitude, we calculated the individual mean response bias per factor level at the corresponding motion direction difference (current trial versus previous trial) that was closest to the obtained peak of the fitted curve. For example, if the fitting procedure yielded a w parameter that indicated a curve peak at 22°, we obtained the individual response errors for 20° and −20°, inverted the response error for the negative distance and then averaged the two values. For the width of the curve, we calculated individual FWHM estimates. Therefore we first smoothed the individual mean response biases across all motion direction difference per condition using locally weighted non-parametric regression fitting (LOESS, implemented as the fLOESS MATLAB function by Marsh, 2016; analog to e.g. Bliss, Sun, & D’Esposito, 2017). We then calculated the FWHM of the smoothed function corresponding to its individual maximum peak between 0 and 60°. If the individual maximum was negative, FWHM was set to 0°. Based on those estimated individual amplitudes and FWHMs we calculated an estimated Cohen’s d as effect size in the following way for the comparison against zero:

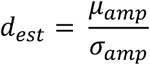

where μ_amp_ is the mean of the individual amplitude estimates and σ_amp_ their standard deviation. For the amplitude contrasts between factor levels, we calculated an estimated Cohen’s d as:

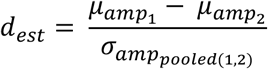

with 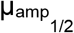 as the mean amplitude for the first and second factor level, respectively and 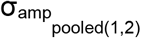 the pooled standard deviation. To calculate the effect sizes for the width contrasts between factor levels, the same formula was used as for amplitude contrasts, but with 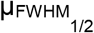 and 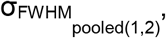 respectively.

#### Software

All Analysis were performed with MATLAB 2018a and the following toolboxes/functions: Circular Statistics Toolbox (Berens, 2009), fLOESS (Marsh, 2016), BADS (Acerbi & Ma, 2017), EzyFit (Moisy, 2016).

#### Code availability

Code is available from the authors upon request.

#### Data availability

Data are available via https://osf.io/azpwy and will be made publicly accessible upon publication.

### Results

#### Serial dependence across trials

In line with previous findings (Bliss et al., 2017; Czoschke et al., 2019; Fischer & Whitney, 2014; Fritsche, Mostert & de Lange, 2017), we observed significant serial dependence across trials even though in this experiment two items were encoded and one was cued for report. Specifically, the reported motion direction of the cued item in the current trial was systematically attracted toward items presented in the previous trial, irrespective of whether the current and previous items shared the same color, serial position or whether the previous item was task-relevant. This attraction followed a DoG-shaped curve with an amplitude parameter of 1.59° (bootstrapped SD: 0.303°, lower 95% of permutations between −1.38° and 0.89°, *p* < *.001*, permutation test (n = 20 participants), *d*_*est*_ = *0.966*, *R*^*2*^ = *.078*). In line with previous studies a maximum attraction was observed for relative small motion direction differences of about 21.44° and a w parameter of 0.033, equaling a width of 34.38° (full width at half maximum, FWHM).

#### Effects of context features

Serial dependence was clearly modulated by the task-relevant context feature, i.e., color (Fig. 2A). We observed a larger serial dependence when the current item had the same color as a stimulus of the previous trial (amplitude = 2.32°, SD = 0.295°, lower 95% of permutations between −1.95° and 1.16°, *p* < *.001*, *d*_*est*_ = *1.359*, *R*^*2*^= *.091*) as compared to when they had different colors (amplitude = 0.87°, SD = 0.646°, lower 95% of permutations between −1.30° and 0.84°, *p* = *.038*, *d*_*est*_ = *0.252*, *R*^*2*^ = *.012*) (amplitude difference = 1.46°, *p* < *.001*, *d*_*est*_ = *0.856,*). Similarly, the observed serial dependence was more broadly tuned when the current item had the same color as a previous stimulus (FWHM = 37.45°) as compared to when they had different colors (FWHM = 28.59°) (w difference: −0.010, equals FWHM difference: 8.86°, *p* = *.047*, *d*_*est*_ = *0.727*).

**Figure 2.**
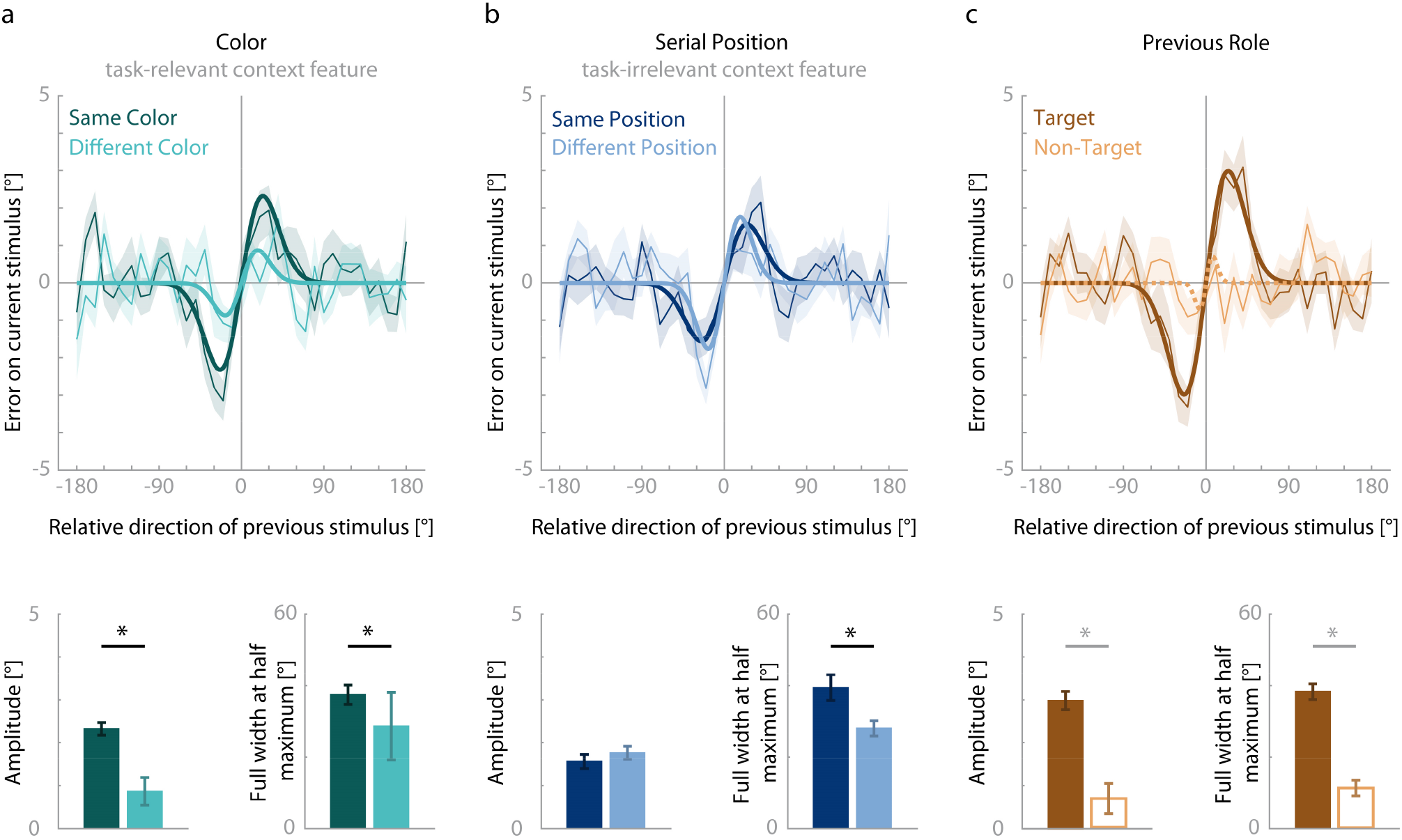
Results of Experiment 1. The response errors (ordinate) are shown as a function of the motion direction difference (abscissa) between an item of the previous trial and the target of the current trial. Positive values on the abscissa indicate that the target direction was shifted counterclockwise relative to an item of the previous trial. Positive values on the ordinate indicate that the response direction deviated clockwise from the true target direction. Serial dependence was revealed by the group averages of response errors (thin lines), with the corresponding shaded regions depicting the standard error of the group mean. A derivative of Gaussian (DoG, model fit shown as bold lines) was fitted to the response errors to estimate the systematic response bias. Solid lines indicate a significant bias, whereas dashed lines depict a non-significant and therefore non-interpretable bias. Bar plots depict the amplitudes and full widths at half maxima (FWHM) with error bars reflecting one SD of a bootstrapped distribution of the parameters. Asterisks indicate significant differences between compared factor levels. a) Both amplitude and width of serial dependence were greater between items with the same color than between items with different colors. b) The width of serial dependence was greater between items with the same serial position than between items with different serial positions. c) A significant serial dependence was observed from a target of the previous trial, but not from a previous non-target. See the online article for the color version of this figure.

In contrast, serial position modulated serial dependence only partially (Fig. 2B). The strength of serial dependence was comparable when the current stimulus was presented at the same serial position (amplitude = 1.56°, SD = 0.331°, lower 95% of permutations between −1.60° and 0.89°, *p* < *.001*, *d*_*est*_ = *0.520*, *R*^*2*^= *.052*) as compared to when they were presented at different serial positions (amplitude = 1.76°, SD = 0.303°, lower 95% of permutations between −1.43° and 0.94°, *p* < *.001*, *d*_*est*_ = *1.178*, *R*^*2*^ = *.046*) (amplitude difference = −0.20°, *p* = *.239*, *d*_*est*_ = *-0.328*). However, we observed a more broadly tuned serial dependence when the current stimulus was presented at the same serial position (FWHM = 39.45°) as compared to when they were presented at different serial positions (FWHM = 28.03°) (w difference: −0.012, equals FWHM difference: 11.42°, *p* = *.011*, *d*_*est*_ = *0.266*).

#### Effects of previous role

Serial dependence was also influenced by the role of an item in the previous trial (Fig. 2C). We observed a serial dependence from previous targets (amplitude = 2.99°, SD = 0.428°, lower 95% of permutations between −2.80° and 1.40°, *p* < *.001*, *d*_*est*_ = *1.351*, *R*^*2*^ = *.140*; FWHM = 38.30°), but no significant attractive bias from previous non-targets (amplitude = 0.71°, SD = 0.699°, lower 95% of permutations between −1.23° and 0.72°, *p* = *.0519*, *d*_*est*_ = *0.358*, *R*^*2*^ = *.003;* FWHM = 11.33°). Therefore, the parameters of the curve fitting analysis are not interpretable for the effect from the non-targets. However, since the permutation test was close to being significant, we calculated the test between the two factors, which showed a stronger and more broadly tuned serial dependence from targets then from non-targets (amplitude difference = 2.28°, *p* < *.001*, *d*_*est*_ = *1.042*; w difference: −0.070, equals FWHM difference: 26.97°, *p* < *.001*, *d*_*est*_ = *1.359*), but emphasize that this result is only restricted interpretable.

#### Interaction effects of task-relevant and task-irrelevant context features and previous role

In addition to the computed contrasts, we examined possible interactions between those three contrasts. First, we examined the interaction of color and serial position. While there was no interaction for amplitude (difference of differences in a: 0.07°, *p* = *.919*, *d*_*est*_ = *0.072*), we observed a significant interaction for width (difference of differences in w: 0.024, equals FWHM difference: −10.77°, *p* = *.041*, *d*_*est*_ = *-0.077*). The width difference between the same versus different serial positions was more pronounced when the items had different colors (w difference: −0.0304, equals FWHM difference: 18.64°) than when they had the same color (w difference: −0.0066, equals FWHM difference: 7.87°). This effect was driven by the difference between items with the same (FWHM = 33.04°) compared with different colors (FWHM = 18.64°) when both had a different serial position. For stimuli of the same serial position, the tuning widths were quite similar for items of the same (FWHM = 40.91°) and different color (FWHM = 37.28°). As only the target showed a significant serial dependence and the influence of the non-target was not clear, we computed the interactions for the other two factors with the previous role of an item, but emphasize that they are only restricted interpretable. We observed a significant interaction between serial position and previous role for width (difference of differences in w: −0.037, equals FWHM difference: 10.80°, *p* = *.022*, *d*_*est*_ = *0.121*). While for targets only, the effect of serial position on width had the same direction as the overall observed effect (w difference: −0.004, equals FWHM difference: 5.16°), for non-targets the effects was reversed (w difference: −0.033, equals FWHM difference: −5.63°), as serial dependence was broader for items with different serial positions. However it should be noted that the width obtained for non-targets with the same serial position was fitted as w = 0.1, which equals the upper bound of the fitting procedure for the w parameter. This indicates that no plausible fit was obtained for this subset of trials, as was already evident in the non-significance of the fit to the non-target trials (see above). We therefore restrain from interpreting this interaction. The interaction of widths was not significant for previous role and color (*p* = *.153*) as well as the interactions of amplitude that involved previous role (*minimum p* = *.311*).

### Results summary

Experiment 1 showed that context features clearly influence serial dependence of object content, but to a different extent. Specifically, both the amplitude and the tuning width of serial dependence were enhanced between items of the same color in comparison to items of different colors. In contrast, while there was no effect of serial position on amplitude, the tuning width of the bias was broader between items with same compared with different serial positions. While the amplitude reflects the strength of the bias, the tuning width describes how similar two consecutive items had to be in order to produce a bias between them. Therefore, while serial position did not affect the strength of the bias, it modulated how similar two items had to be in order to observe a bias between them. Importantly, in this experiment color was the task-relevant context feature, which served for cueing, whereas serial position was task-irrelevant. However, the observed differences in serial dependence between these context features cannot be unambiguously attributed to task relevance, because both features may also differ in salience. Color is a salient feature inherent in the visual appearance of an item. On the other hand, serial position defines the temporal structure of a trial and is inherent in the encoding of the stimuli even if it is task-irrelevant. To test whether the modulation by context congruence was determined by whether the context feature was task-relevant, we swapped the task relevance of both features in Experiment 2.

Experiment 1 also revealed that a significant attractive bias was only observed towards target items of the previous trial. This result is consistent with our recent study that also found serial dependence in a WM paradigm only towards the item that was cued for report (Czoschke et al., 2019). This indicates that when multiple objects were encoded into WM within one trial, only the object that was internally selected for response caused serial dependence. However, since there was a trend towards a positive serial dependence produced by the non-target, this conclusion has to be treated with caution.

To overcome the limitations of Experiment 1 we performed Experiment 2 in which we used serial position as task-relevant context feature. We also eliminated any association between serial position and stimulus color by allowing two items within a trial to have the same color. In addition, to obtain more conclusive evidence about the potential serial dependence on non-targets, we substantially increased the number of participants.

## Experiment 2: Serial Order Cueing

### Methods

#### Subjects

Fifty-one subjects who were recruited from the Goethe-University Frankfurt and the Fresenius University of Applied Sciences Frankfurt participated in Experiment 2, none of whom had participated in Experiment 1. All subjects reported normal or corrected-to-normal vision. Two subjects were excluded from the final analysis due to poor task performance (SD of report error > 3 SDs of the sample mean). We thus included 49 subjects (19 male), aged between 18 and 33 years (mean: 23.8 years). All subjects gave informed consent and were compensated with € 10/h or course credit. The study was approved by the Ethics Committee of the Medical Faculty of the Goethe-University Frankfurt am Main and therefore complied with their ethical regulations.

#### Procedure and stimuli

The procedure and stimuli used equaled the ones described for Experiment 1. There were only two differences. First, cueing was now based on the encoding position of the stimuli instead of their color. A number cue replaced the fixation square to indicate which one of the presented directions had to be reported (Figure 1b). To ensure that stimulus color yielded no information about the encoding position of the stimulus, as a second change the stimuli presented in one trial could now have either the same or different colors. This resulted in four possible color combinations within a trial (red-red, red-green, green-green, green-red). Each of those color combinations occurred equally often. In half of the trials, the first presented stimulus had to be reported, and in the other half the second one, which was balanced over the different color and direction difference combinations. Three different monitors with a 4:3 display ratio were used, two running with 60 Hz and one with 75 Hz refresh rate, but with all stimulus parameters kept constant. Subjects were again seated at a viewing distance of approximately 50 cm.

#### Analysis

All conducted analysis steps were the same as in Experiment 1.

### Results

#### Serial dependence across trials

When two items were encoded into WM and one of them was cued for report via serial position we observed a significant serial dependence across trials with an amplitude of 2.00° (SD = 0.200°, lower 95% of permutations between −0.93° and 0.57°, *p* < *.001*, *d*_*est*_ = *1.123*, permutation test (n = 49 participants), *R*^*2*^ = *.118*) and a FWHM of 34.63°, with the peak of the curve located at 21.44°.

#### Effects of context features

Serial dependence was clearly modulated by the task-relevant context feature, i.e., serial position (Fig. 3B). The strength of serial dependence was enhanced when the current stimulus was presented at the same serial position as a previous one (amplitude = 2.33°, SD = 0.204°, lower 95% of permutations between −1.15° and 0.71°, *p* < *.001*, *d*_*est*_ = *1.243*, *R*^*2*^ = *.106*) as compared to when they were presented at different serial positions (amplitude = 1.82°, SD = 0.200°, lower 95% of permutations between −0.87° and 0.56°, *p* < *.001*, *d*_*est*_ = *0.881*, *R*^*2*^ = *.045*) (amplitude difference = .52°, *p* = *.009*, *d*_*est*_ = *0.387*). Additionally, we observed a more broadly tuned serial dependence between items at the same serial position (FWHM = 44.60°) compared with different serial positions (FWHM = 26.04°) (w difference: −0.018, equals FWHM difference: 18.56°, *p* < *.001*, *d*_*est*_ = *0.840*).

**Figure 3.**
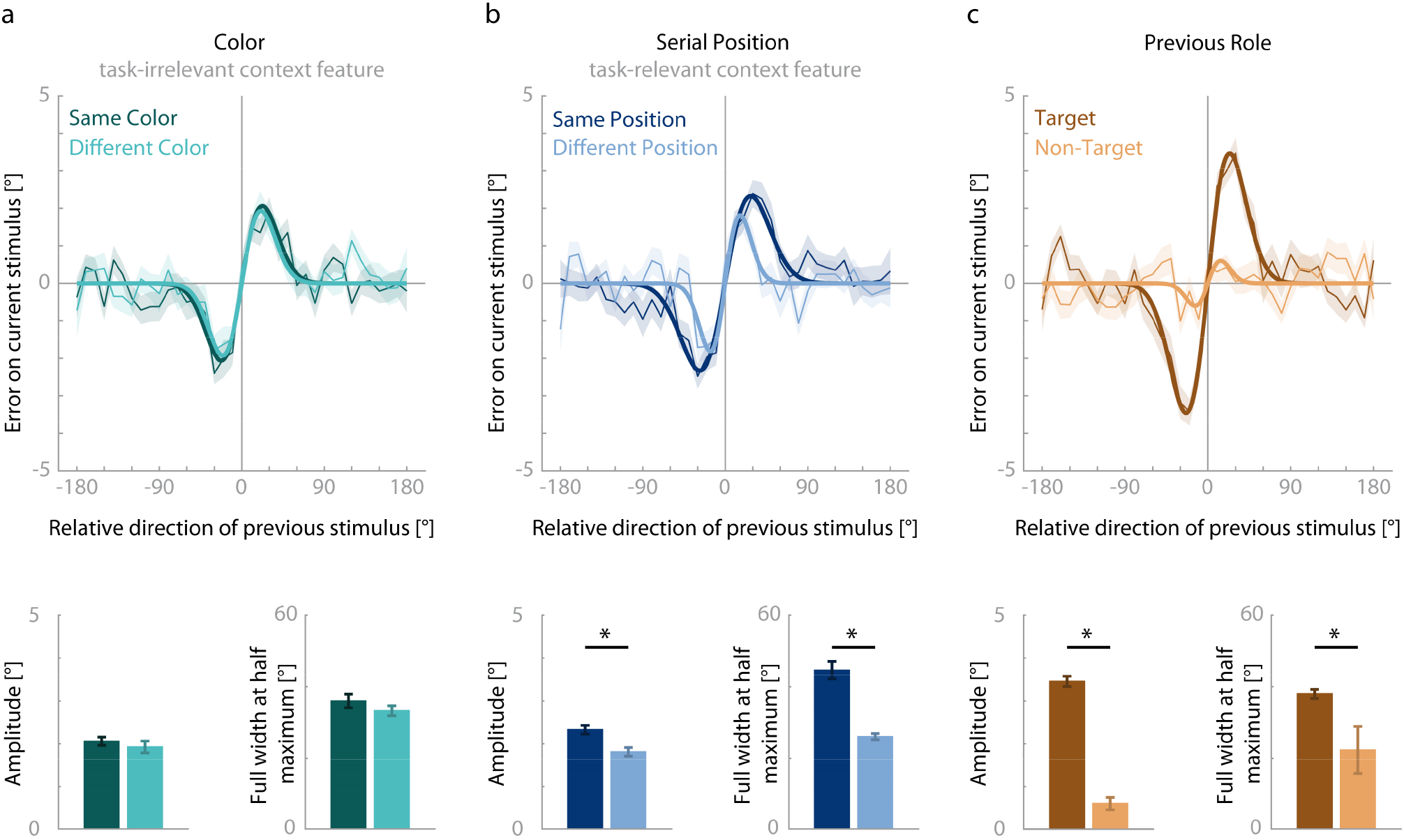
Results of Experiment 2. Serial dependence (ordinate) was shown as a function of the motion direction difference (abscissa) between an item of the previous trial and the target of the current trial. For details see Figure 1 and Methods. a) Serial dependence did not differ significantly between items with the same color and different colors. b) Both amplitude and width of serial dependence were greater between items with the same serial position than between items with different serial positions. c) Both amplitude and width of serial dependence were greater for previous targets than non-targets.

In contrast, the task-irrelevant context feature color did not modulate serial dependence (Fig. 3A). Specifically, we observed a comparable strength of serial dependence when the current item had the same color as a previous stimulus (amplitude = 2.06°, SD = 0.195°, lower 95% of permutations between −1.19° and 0.65°, *p* < *.001*, *d*_*est*_ = *0.674*, *R*^*2*^ = *.076*) as when they had different colors (amplitude = 1.93°, SD = 0.275°, lower 95% of permutations between −1.33° and 0.63°, *p* < *.001*, *d*_*est*_ = *0.812*, *R*^*2*^ = *.057*) (amplitude difference = 0.13°, *p* = *.347*, *d*_*est*_ = *-0.065*). The same was true for tuning widths (same color: FWHM = 36.00°, different colors: FWHM = 33.27°, w difference = −0.0003, equals FWHM difference: 2.73°, *p* = *.228*, *d*_*est*_ < *0.001*).

#### Effects of previous role

Experiment 2 confirmed that the role of an item in the previous trial, i.e. whether it was a target or not, strongly modulated serial dependence (Fig. 3C). Again, we observed a serial dependence from previous targets (amplitude = 3.46°, SD = 0.240°, lower 95% of permutations between −1.59° and 0.92°, *p* < *.001*, *d*_*est*_ = *1.654*, *R*^*2*^ = *.176*). However, increasing the number of subjects revealed an attractive bias from previous non-targets, too (amplitude = 0.60°, SD = 0.297°, lower 95% of permutations between −0.62° and 0.44°, *p* = *.009*, *d*_*est*_ = *0.421*, *R*^*2*^ = *.005*). Importantly, the bias from a previous target was stronger than from a previous non-target (amplitude difference: 2.86°, *p* < *.001*, *d*_*est*_ = *1.192*). Furthermore, the tuning width of the serial dependence on previous targets (FWHM = 37.98°) was broader than on previous non-targets (FWHM = 22.28°) (w difference = −0.021, equals FWHM difference: 15.70°, *p* < *.001*, *d*_*est*_ = *1.070*).

#### Interaction effects of task-relevant and task-irrelevant context features and previous role

We also examined possible interactions between the three contrasts. None of the computed interactions was significant. For the amplitudes, we observed one weak trend (*p* = *.111*, *d*_*est*_ = *-0.140*) suggesting that the observed amplitude enhancement between items with the same serial position was more prominent when the previous item was a non-target (amplitude difference: 1.12°) than when it was a target (amplitude difference: 0.48°). For the non-target items, position incongruence in fact might even lead to a reversal of the observed bias (position congruent non-targets: 0.61° amplitude, position incongruent non-targets: −0.51° amplitude). All other interactions were far from reaching significance both for amplitude (*minimum p* = *.610*) and width (*minimum p* = *.464*).

### Results summary

Experiment 2 aimed at resolving two open questions: first, whether task-relevant and task-irrelevant context features differentially modulate serial dependence of object content, and second, whether serial dependence also occurs for non-targets from previous trials. We found that color that served as task-relevant context feature in Experiment 1 but was task-irrelevant in Experiment 2 did not modulate serial dependence. Similarly, task-relevant serial position in Experiment 2 affected the amplitude of serial dependence, which was not the case in Experiment 1 where serial position was task-irrelevant. However, in contrast to color, serial position affected the tuning width of serial dependence regardless of whether it was task-relevant or not. Regarding the effect of the previous role on serial dependence, Experiment 2 with an increased number of subjects revealed an attractive bias also from previous non-targets. However, both amplitude and tuning width of serial dependence was notably smaller for previous non-targets than targets.

### General Discussion

Current models assume that objects in WM consist of bound content and context features that are established anew in each memory episode. Recent studies have shown, however, that the content feature of a single object encoded into WM was selectively attracted toward a similar content feature of a past object representation. This phenomenon, termed serial dependence, has attracted much research interest (Kiyonaga et al., 2017). But, as we typically hold several objects in WM, it has remained unclear whether multiple objects interact across memory episodes. To answer this question, we conducted two experiments in which participants memorized two sequentially presented motion directions (S1 and S2) that differed in color. After a brief delay, either a color cue (red or green; Experiment 1) or a serial position cue (first or second stimulus; Experiment 2) indicated which motion direction stimulus to report. We could thus assess the impact of three factors on serial dependence: congruence of context features across trials (color or serial position), task relevance of context features, and the role of the object in the previous trial. We found that all these factors support serial dependence in situations where several objects are encoded into WM. Specifically, we observed a stronger serial dependence between items that shared the same task-relevant context features across trials. Moreover, serial position partly facilitated serial dependence even when it was the task-irrelevant context feature, whereas task-irrelevant color had no such effect. Third, regardless of the context feature, the attractive bias was stronger toward the target item in the previous trial. Together, our results show that serial dependence based on content-similarity is enhanced between objects that share the same task-relevant context features and that are internally selected as target objects.

In most previous studies on serial dependence between trials, only one item per trial had to be encoded into WM. In contrast, the present study found that serial dependence also occurs when two items per trial were attended and encoded into WM, replicating recent results from our laboratory (Czoschke et al., 2019). Importantly, we also replicated the finding that serial dependence across trials is particularly pronounced for target items that were retro-cued for report in the previous trial. Retro-cueing implies that targets were internally selected into the focus of attention. In contrast, previous studies have found enhanced serial dependence for pre-cued targets, i.e., as a result of externally manipulating the attentional selection and encoding (Fischer & Whitney, 2014). Taken together, these findings show that both external and internal attention strongly promote serial dependence. Interestingly, the present Experiment 2 with an increased number of subjects revealed serial dependence also for non-targets, albeit with a clearly reduced amplitude compared to targets. As both targets and non-targets from the previous trial were irrelevant for the current trial, this result indicates that attentional prioritization in the previous trial strongly modulates the magnitude of serial dependence, without being a necessary precondition for its occurrence.

Besides the influence of attention, another key property of serial dependence is that it operates selectively between objects with similar contents, as reflected by the stereotypical tuning profile observed for a range of different content features like orientation (Fischer & Whitney, 2014), faces (Liberman et al., 2014), spatial position (Bliss et al., 2017), ensemble representations (Manassi et al., 2017) or motion direction (Czoschke et al., 2019). Based on this property, Fischer and Whitney (2014) suggested that serial dependence reduces small differences between consecutive content features to support the impression of a coherent environment. The novel finding of the present study was that, in addition to content similarity, context features also leave traces in WM and thus help to relate corresponding objects across memory episodes. Specifically, we observed a stronger and more broadly tuned serial dependence between motion directions with the same color (Experiment or serial position (Experiment 2). Previous studies using only one stimulus per trial have yielded contradictory results concerning the role of context features like spatial position for serial dependence (Fischer & Whitney, 2014; Fritsche et al., 2017). The present design requiring the selective report of one out of two items per trial should have increased the binding between content and context features, thus enhancing the influence of context on serial dependence. This finding corroborates the assumption that serial dependence indicates a continuity field in a changing environment that promotes stability of object representations over time.

The observed modulation of serial dependence by both content similarity and context congruence suggests that single features of an object are represented in WM as bound together to some degree. This is in line with frameworks that assume a WM organization with integrated multi-feature objects (Brady et al., 2011; Luck & Vogel, 1997; Oberauer & Lin, 2017). The definition of a WM item as a combination of content and context features corresponds closely to the concept of object files (Kahneman, Treisman, & Gibbs, 1992; Treisman, 1986). An object file contains the different features of an object that form a temporary representation, enabling us to track an object over time. The object identity should remain stable and immune against small changes in object appearance, caused, e.g., by movement or changing lighting conditions. While priming studies have shown that object features are discriminated faster for previously presented objects (Noles, Scholl, & Mitroff, 2005), the mechanism enabling the tracking of objects over time has remained unknown. Fischer and Whitney (2014) have proposed serial dependence as a mechanism underlying the continuity of objects. However, across memory episodes with multiple relevant objects, serial dependence would serve temporal integration only if it operated in an object-file fashion by relating corresponding bundles of content and context features. Our findings show that context congruence of objects clearly promotes serial dependence, thus supporting the interpretation that serial dependence is a mechanism suited for temporal integration of object representations across time.

While task-relevant context features consistently enhanced serial dependence, the effect of the task-irrelevant context feature differed between experiments. In Experiment 1, we found that task-irrelevant serial position changed the tuning width of serial dependence. In contrast, in Experiment 2 there was no such effect for task-irrelevant color. One explanation might be that serial position is more automatically integrated into an object representation than color, even though the latter one is a more salient visual feature. This relates to the importance of spatiotemporal information for the definition of an object (Treisman, 1986). Furthermore, it reflects the importance of serial position as a context feature regardless its task-relevance. The sequential order of events, i.e., stimulus presentations, determines the serial position of an item and thus defines the temporal structure of a trial. Therefore, temporal information might be of crucial relevance for item representations, particularly in the case of sequentially presented items at the same spatial position (Schneegans & Bays, 2018). Taken together, future studies should investigate the role of temporal position for WM representations in greater depth.

Furthermore, we observed different effects of task-relevance on the amplitude of the bias for both context features. For color, our results indicate that serial dependence was suppressed between differently colored items when it was task-relevant (amplitude = 0.87°) compared to when it was task-irrelevant (amplitude = 1.93°) whereas it was quite similar for items of the same color when it was task-relevant or task-irrelevant (amplitudes of 2.32° and 2.06°, respectively). For serial position in contrast, task-relevance enhanced the amplitude of serial dependence between items with the same serial position (amplitude = 2.33°) in contrast to when it was task-irrelevant (amplitude = 1.56°) and was similar for items with different serial positions when it was task-relevant or task-irrelevant (amplitudes of 1.82° and 1.76°, respectively). This indicates that task-relevance enhanced the amplitude between items with the same serial position whereas it suppressed the amplitude between items of different colors. This further strengthens the interpretation that color and serial position might be two qualitatively different context features.

A large body of research on WM has shown that when a probe did not form part of the currently memorized set but of the set presented on the previous trial, reaction times were longer and recognition accuracy was reduced (see Jonides & Nee, 2006, for an overview). This phenomenon is termed proactive interference and has been investigated most commonly using verbal stimuli (e.g., Jonides, Smith, Marshuetz, Koeppe & Reuter-Lorenz, 1998; Keppel & Underwood, 1962). Proactive interference has been observed also for visual features like colors or shapes. Here it was particularly pronounced when past and current items were presented at the same spatial location (Makovski & Jiang, 2008). Similarly, in serial recall tasks participants often incorrectly reported items from previous lists (so-called intrusions) at the same recall position across trials (e.g., Henson, 1999). Proactive interference and serial dependence both describe effects of previous WM episodes on current ones, which has led to the assumption that both could arise from the same underlying mechanism (Kiyonaga et al., 2017). Our results demonstrate that serial dependence is supported by corresponding context features across trials. As the same is true for proactive interference (Henson, 1999; Makovski & Jiang, 2008), this supports the possibility that serial dependence and proactive interference reflect the same mechanism. On the other hand, this hypothesis is challenged by important differences between both phenomena. Proactive interference effects arise because an item from a previous trial is mistakenly assigned to the item set of the current trial. This indicates that the binding between the item and its trial context was incorrect. In contrast, serial dependence describes an integration of a previous into a current content feature of an object that is promoted by content similarity and context congruence, rather than the erroneous recall of a previous object. Furthermore, in most studies on proactive interference, the observed effects stemmed from previously encoded but not tested items, i.e., non-targets. Bäuml and Kliegl (2013) even showed that testing items of a previous list eliminated proactive interference on a subsequent list. On the other hand, serial dependence mainly arises from the target of a previous trial, as shown by our results together with previous ones (Czoschke et al., 2019; Fischer & Whitney, 2014). Moreover, proactive interference is usually interpreted as a malfunction that has to be overcome whereas serial dependence is considered a beneficial temporal smoothing mechanism. Our results support the idea that the mechanism underlying serial dependence is a beneficial one, because it relates objects over time by reducing small differences between their representations. While this would be useful in natural environments, it may lead to harmful outcomes in artificial settings, leading to systematic errors or reduced performance in studies of serial dependence or proactive interference, respectively. Notably, proactive interference has been found to correlate negatively with both WM capacity (Mecklinger, Weber, Gunter, & Engle, 2003) and intelligence (Braver, Gray, & Burgess, 2007). If interference effects result from a generally beneficial mechanism, individuals with higher WM capacity or fluid intelligence might be better at strategically controlling this mechanism. Taken together, proactive interference and serial dependence differ with regard to their conceptual explanation and whether they arise from previous targets or non-targets, but both describe an influence from past memory events on current ones, which can be modulated by context congruence of objects across time. Therefore, more research is needed to disentangle if and to which degree those phenomena stem from the same underlying mechanism.

Until now, the processing stage at which serial dependence occurs has remained unclear. While some studies have suggested a perceptual stage (Cicchini, Mikellidou, & Burr, 2017; Fischer & Whitney, 2014; Fornaciai & Park, 2018; Manassi et al., 2017; St. John-Saaltink, Kok, Lau, & de Lange, 2016), others have provided evidence for a memory- or decision-related process (Bliss et al., 2017; Fritsche et al., 2017; Papadimitriou, Ferdoash, & Snyder, 2015; Pascucci et al., 2019). The present findings support the hypothesis of serial dependence as a memory-based or decisional mechanism. We observed that the impact of both investigated context features on serial dependence relied heavily on task-relevance of the features. Furthermore, the purely visual context feature, i.e. color, relied more strongly on task-relevance than serial position. Importantly, the latter feature is not inherent in the visual presentation of a stimulus but only arises within the context of a trial. Taken together, these observations argue against an exclusively perceptual basis of serial dependence. Future investigations of the neural underpinnings of this mechanism could elucidate the processing stage at which serial dependence occurs.

Our study showed that representations in WM are biased towards previous representations, specifically to those that were targets and had corresponding task-relevant context features. This provides a new insight into the organization of object processing. Apparently, the binding of content and context features is not completely erased after a memory episode, but to some extent is carried over to subsequent episodes pointing toward a mechanism that selectively integrates corresponding multi-feature object representations over time.

## Acknowledgements

This study was supported by the German Academic Scholarship Foundation (PhD Scholarship awarded to C.F.). We thank Plamina Dimanova and Alina Rebitzky for their help in data collection as well as Julia Krebs for help with piloting.

